# Cognitive Entities Underpinning Task Episodes

**DOI:** 10.1101/216796

**Authors:** Ausaf A Farooqui, Tom Manly

## Abstract

Accounts of hierarchical cognition suggest that extended task episodes as one task entity and not individually execute their component acts. Such hierarchical execution is frequently thought to occur by first instantiating a sequence representation in working memory that then controls the identity and sequence of component steps. In contrast, we evidence hierarchical execution of extended behavior in situations where the identity and sequence of component steps was unknown and not linked to any sequence representation. Participants executed unpredictable trials wherein, depending on the margin color, they could either choose the smaller value or the font of the two numbers. Crucially, they were biased into construing a recurring instance of three or five trials as one task episode. Behavioral signs of hierarchy, identical to those seen previously with memorized sequence execution, were seen - High trial 1 RT, higher trial 1 RT before longer task episodes, and absence of trial level switch cost specifically across episode boundaries. This showed that hierarchical task sequence representations are not a requisite for hierarchical organization of cognition. Merely construing behavior to be executed as a task episode is enough for the accompanying cognition to be hierarchically organized. Task episodes are executed through the intermediation of episode related programs that are assembled at the beginning and control and organize cognition across time.

## Introduction

Goal-directed behavior frequently consists of temporally extended actions and task entities (task episodes, sub-episodes) that consist of a sequence of steps (e.g. subtasks, smaller acts and events etc.) but are nonetheless executed as one entity. The action of pouring milk in a cup is selected and instantiated as one action even though it consists of a coordinated sequence of numerous smaller motor acts; likewise, the various components of checking email at different levels of detail (‘open browser’, ‘move cursor’, ‘click’) are executed as one entity (e.g. Botvinick, Niv & Barto 2009; Dezfouli, Lingawi, & Balleine, 2014; Logan, 1988). In what sense are the sequence of acts corresponding to such task episodes executed as *one* entity is unclear. More so, because what constitutes *a* task is often determined by the doer’s construal and not by the physical environment (Vallacher & Wegner, 1989). The same breakfast, for example, can be prepared as one (‘prepare breakfast’), two (‘prepare tea’ and ‘prepare toast’) or four task episodes (‘boil water’, ‘brew tea’, ‘toast bread’, ‘spread butter).

The experiments presented here examine the cognitive basis of extended tasks. We use the term *task episodes* because these typically are temporally extended periods, with a defined beginning and an end, of focused purposive behavior during which a sequence of constituent steps are executed. Here, while participants executed continuous trials of an experimental session, task-irrelevant cues were used to bias them towards viewing each recurring period consisting of three or five trials as *one* task episode. This method allowed us to study factors related to task-episodes without confounding them with factors pertaining to task rules, working memory, attention, action selection etc. While we use the phrase ‘task-episode’ in this paper, we do not insist that in the task hierarchy these are tasks as opposed to sub-tasks of a larger task. Our concern is that irrespective of the hierarchical level, the sequence of trials be construed and executed as one entity.

### Hierarchical Organization of Actions

When a sequence of smaller acts is executed as one action, the larger action is necessarily hierarchical. While the presence of hierarchy in behavior has long been recognized (e.g. Lashley, 1951; Miller, Galanter & Pribram, 1960), its details are still being explored. Notably, hierarchical actions have been characterized in situations that are predictable, where the knowledge (both procedural and declarative) related to the identity and sequence of component steps is accessible as a *single entity* described variously as schemas, scripts, frames, plans (Minsky, 1975; Miller et al. 1960; Norman & Shallice, 1986; Rumelhart & Norman, 1985; Schank & Abelson, 1977). This entity is thought to form the higher-level of the cognitive hierarchical organization and controls the identity and sequence of lower-level steps. For example, the schema about preparing coffee may consist of ‘add coffee from packet’, ‘add sugar from bowl’, ‘add milk from carton’, each of which may have their own sub-schemas. The main schema may be active across time and sequentially select the lower level schemas.

Experimentally, when participants generate a memorized sequence of motor acts, recall a memorized sequence, or execute a memorized list of task items – the corresponding extended behavior shows three characteristic signs that suggest its execution as one hierarchical cognitive entity (e.g. Anderson et al. 1998; Kahana & Jacobs, 2000; Rosenbaum, Kenny & Derr, 1983; Schneider & Logan, 2006). First, item 1 reaction time (RT) tends to be the longest. Second, this item 1 RT is frequently longer for longer/more complex sequences, e.g. it takes longer to start speaking longer words, execute task lists with more item-level switches. Third, the behavioral cost of switching (or benefits of repeating) tends to be absent when component items switch (or repeat) across sequence boundaries. Hence when sequence like AABB (where A and B stand for different task items, e.g. color and shape judgements, to be executed on individual trials) is repeatedly executed (AABB-AABB…) the first position is a switch from the last executed item. In contrast, during repeated execution of a sequence like ABBA (ABBA-ABBA…) the first position is a repeat from the previous item. Still the step 1 RT or error rate for AABB is not higher than for ABBA (Lien & Ruthruff, 2004; Schneider & Logan, 2006, 2015).

These three signs of hierarchy have frequently been interpreted as resulting from the dynamics of recall and instantiation of a hierarchical and propositional task-sequence representation in working memory that controls the identity and sequence of component steps (e.g. Lien & Ruthruff, 2004; Mayr, 2009; Perlman et al. 2010; Schneider & Logan, 2006 and 2015). Beginning to execute the sequence requires instantiating the sequence representation in working memory, hence the longer item 1 RT, and longer sequences take longer to instantiate causing longer item 1 RTs for longer sequences. Processes instantiating a hierarchical sequence structure in working memory remove the switch cost related to component items (Schneider & Logan, 2006 and 2015).

However, it is also possible that these signs of hierarchical cognition result not from the dynamics of propositional representation of the task sequence in working memory but from the execution of extended behavior as one entity. When participants are given a task sequence to execute, it is plausible that they will conceive of the corresponding behavior as *one* task-episode and execute it as one entity. In this scenario, the higher trial 1 RT may be related to preparing for the entire episode before executing the first item. More complex/longer episodes may take longer to prepare, hence higher trial 1 RT during more complex sequences. Furthermore, this will also lead to absence of component item related switch costs at the beginning of task episodes.

Switch costs are known to be affected by doer’s notion of task (Dreisbach, Goschke & Haider, 2006, 2007; Dreisbach & Haider, 2008; Schneider & Logan, 2014). In a set of experiments by Dreisbach et al. word stimuli appeared in different colored fonts across trials with each color cueing a specific task (e.g. green: animal-nonanimal judgement task on the word, red: consonant-vowel judgement task on the first letter of the word). One group of participants (the ‘two-task’ group) was informed of the two tasks represented by the color-task mappings. This group therefore construed the two colors as cueing two different *tasks.* In contrast, the other group (the ‘stimulus-response’ group) memorized all the stimulus-response mappings. Switch cost associated with change of font color was present in the ‘two-task’ group but not in the ‘stimulus-response’ group, even though the two groups executed trials that were identical in their requirements.

When a sequence of trials is conceived as one task episode, as happens during task-sequence execution, the task identities of individual trials will get subsumed by that of the larger episode. The trials become steps or subtasks of the construed task-episode. Cognitive entities related to individual trials will be at a hierarchically lower level than those related to the episode. In hierarchical relations a change at a higher level will necessarily cause a change at the lower level, but not vice-versa (Broadbent, 1977). Change in task-episode related entities at episode boundaries will necessarily change those related to individual trials, resulting in no advantage of repeating (or cost of switching) a trial item across episode boundaries.

We thus hypothesized that behavioral signs previously documented during the execution of predictable sequences follow primarily from the execution of corresponding behavior as *one* task-episode. This predicted that those signs would be evident whenever a segment of purposive behavior is executed as *one* task episode, even when components of that behavior are unpredictable and do not follow from any memorized sequence representation.

This thesis is supported by our everyday phenomenal experience. We prepare breakfasts, write emails and shop. The very many component acts of these task episodes are executed not as a collection of independent entities, but as parts of one hierarchical task episode. Further, predictable action and task sequences are not the only instances of hierarchical execution. We frequently execute extended behavior hierarchically as *one* task-episode even when the identity and sequence of its component steps is unpredictable and cannot be retrieved as a mnemonic chunk. We can shop without a list in mind, improvise while playing music, speak extemporaneously, edit unfamiliar paper etc. In all such instances, we may not know what all steps we will take during the period, nonetheless, we have a strong phenomenal sense that the behavior during the period is executed as parts of one task-episode and not as a sequence of independent acts. We even reliably identify such task-episodes with unique task identities (Vallacher & Wegner, 1989). If this is not epiphenomenal then there must be specific cognitive means, other than propositional knowledge and representation of component actions and task items, of cohering extended behavior into hierarchical cognitive entities.

One possibility is that cognitive entities following from the doer’s notion of the task and goal being executed subsume and control the execution of behavior. Tasks and goals are intimately related constructs and one cannot be talked of without the other. Goals conceptions are important for the control and execution of actions that lead to their achievement (Anderson, 2014; Gollwitzer & Bargh, 1996; Greenwald, 1972; James, 1890; Jeannerod, 1988; Lewin, 1926; Meyer & Kieras 1997; Prinz, 1987). For these to retroactively control the episode leading to their achievement, some goal-directed cognitive entity has to actively subsume the execution of the task episode (e.g. James, 1890; Kruglanski & Kopetz, 2009). Tasks and goals, therefore, may not just exist as propositional representations, but may be better understood as hierarchical and episodic cognitive entities that subsume cognitive processing across the period conceived as a task-episode that culminates in the completion of the conceived goal.

The presence of such task-episode related entities that go beyond mere task sequence representations was suggested by a neuroimaging experiment (Farooqui, Mitchell, Thompson & Duncan, 2012). Participants were to sequentially search and detect four pre-specified letter targets while viewing a sequence of single letter presentations. Although the four target detections were identical, participants were instructed that the first three searches were part of one subtask, while the fourth one was a separate subtask. If this biasing was successful, participants would conceive the 3rd target detection as completing a subtask episode while the 1st and 2nd target detections would be events lying within this episode that perhaps completed sub-subtasks within this subtask episode. The 4th target detection, on the other hand, completed the main task-episode. Note that the sequential searches were identical and there was nothing structural in them that would distinguish 3rd and 4th target detections from the first two, apart from their position in the conceived task-episode. In fact these searches could be equally well executed if these target detections were conceived as a flat sequence of four searches.

The 3rd target detection completing the conceived subtask episode, elicited greater and more widespread brain activity compared to identical target detections within that episode (i.e. 1st and 2nd); while the 4^th^ target completing the main task-episode elicited maximal activity. This pattern of elicited activity: sub-subtask completion < subtask-episode completion < task-episode completion, could not have resulted from changes in attention, working memory, task rules etc. because the different targets were identical in these terms. The most plausible account was that these activities reflected the conceived organization of the task and resulted from a change in cognitive entities related to the conceived task/subtask episodes. Completion of a lower-level episode elicited limited activity because it dismantled only those cognitive entities that were related to it leaving those related to the higher-level episode intact. Completion of the higher-level episode on the other hand elicited intense and widespread activity because it dismantled cognitive entities at both higher and lower levels of hierarchy.

### Current Study

We predicted that to the extent an extended behavior is *construed* as *a* task episode it will be executed as one cognitive entity, irrespective of the predictability of its components. The three signs of hierarchical cognition seen previously during the recall and execution of memorized sequences will be seen whenever any arbitrary segment of behavior is construed as a task-episode, even when sequence related propositional representations are absent and the corresponding behavior is unpredictable.

Participants executed trials on which one of two items could be executed depending on the color of stimulus margins (blue: choose the lower value between the two numbers; green: choose the smaller font). The item to be executed on any trial was not known beforehand, and the probability of it being repeated or switched across successive trials was the same. Across different experiments a variety of means was used to bias subjects into conceiving a sequence of consecutive trials (or a period of time in experiment 5) as a defined task episode. These included having to additionally count trials in threes (1-2-3) or fives (1-2-3-4-5; experiments 1 and 6), temporal grouping of trials along with an irrelevant countdown (experiments 2, 3 and 4), and the presence of an irrelevant outer margin that stayed on for extended epochs of time and framed the execution of extended series of trials (experiment 5).

If trials of such construed task episodes were executed as parts of a larger cognitive entity and not as independent trials then (1) trial 1 RT will be longest because of the additional time needed to assemble the subsuming task episode related cognitive entity. (2) Trial 1 RT will be longer for longer episodes because episode related cognitive entity related to longer episode will take longer to assemble compared to those related to shorter episodes. (3) Trial rule related switch cost will be absent at position 1, because the dismantling and reassembly of the subsuming episode related entity will refresh lower-level trial rule related configurations.

## Experiment 1

### Methods

Participants were asked to keep a covert count of trials in 3s (1-2-3-1-2-3) or 5s (1-2-3-4-5-1-2-3-4-5) depending on an onscreen ‘steps of 3’ or ‘steps of 5’ instruction at the beginning of each block made of 6-35 trials (Figure 1). This screen remained on until participants pressed the spacebar which began a run of trials. On each trial two numbers differing in value (between 0 to 99) and font size (Arial font 60 or 20) were displayed on each side of the fixation cross. A margin box appeared around, and at the same time as, each number display. Margin color determined the rule relevant on that trial - blue: choose the smaller value, green: choose the smaller font. Participants’ decisions were conveyed via buttons presses that were spatially congruent with their choice (Numpad 1 for left and Numpad 2 for right on a standard QWERTY keyboard number pad). The stimuli remained onscreen until a response was made, and was followed by the next trial after 1 s.

**Figure 1.**
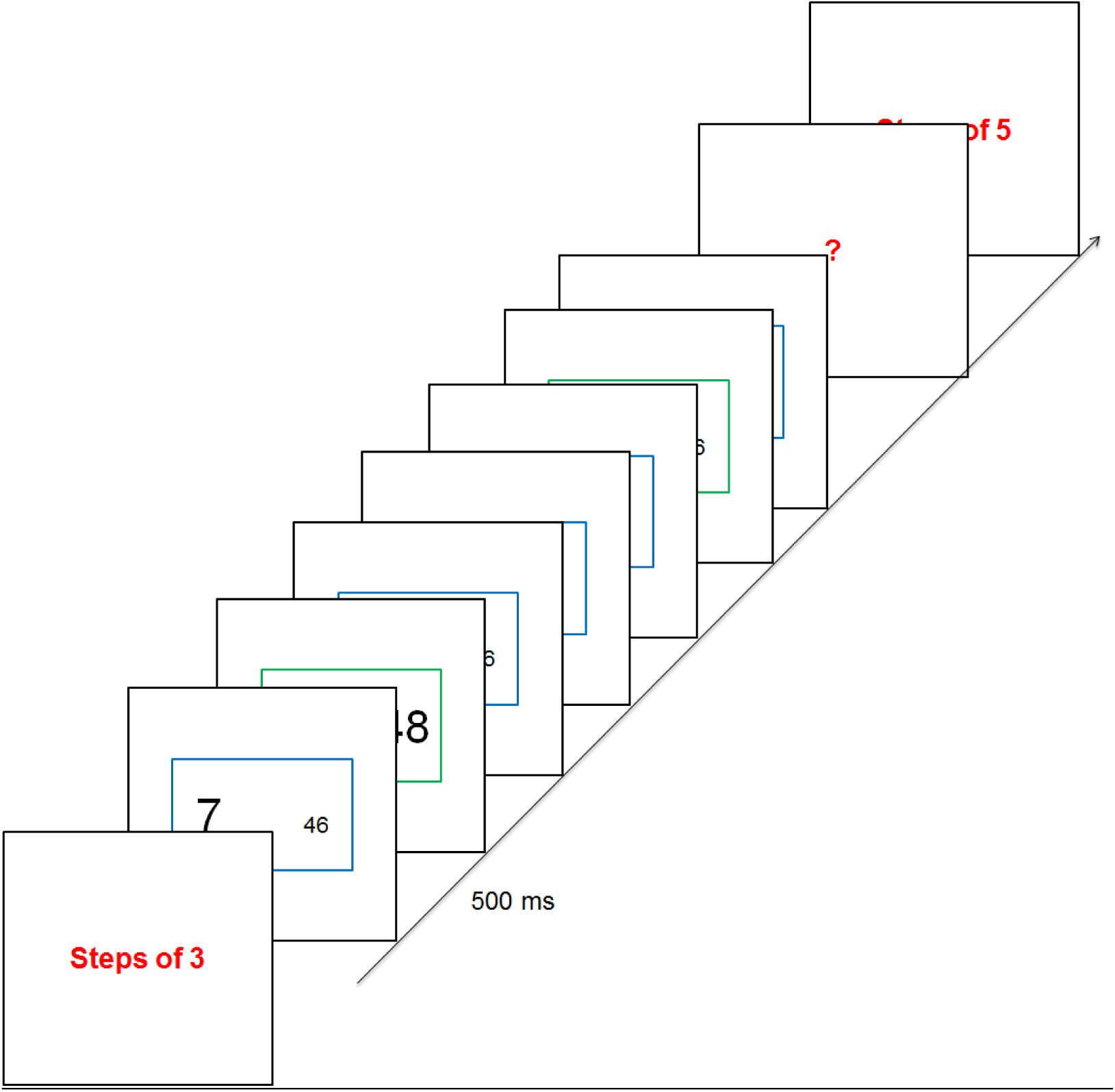
Trial blocks began with an instruction screen stating whether trials were to be executed in steps of 3 or 5. Subsequent trials were to be executed while keeping a covert count (e.g. steps of 3: 1-2-3-1-2-3…). Trial rule was cued by the color of the outer margins – blue: choose the smaller value, green: choose the smaller font. The block would end with a ‘?’, to which participants keyed in the number of the step just executed.

To check whether participants were keeping an accurate 3 or 5-trial count, a probe (“?”) appeared at the end of the block of 6-35 trials. Participants were to key in the step number that they had executed before the probe appeared, e.g. if the probe appeared after 7 trials, the correct response in a 3-trial episode block would be 1 (1-2-3-1-2-3-1). Feedback was given on the accuracy of probe responses (high pitch tone for correct, low pitch tone for incorrect).

This and subsequent experiments were created in Visual Basic.net and run on a Dell computer with an 85 Hz refresh rate monitor kept at a comfortable distance from the participant. The experiment was conducted on an individual basis in a testing room designed to minimize visual and noise distraction. Participants were first given 7 trials of practice on each of the two trial types (value and font judgments). They then completed a 30-trial practice block in which the stimulus margin changed its color randomly signaling the currently relevant rule. They were then told to execute trials while keeping a count in threes. They practiced on two such blocks before proceeding on to the main experimental session. Participants completed 70-120 blocks each consisting of 6-35 trials with 3- and 5-trial blocks being randomly interlaced. Participants were to respond as quickly and as accurately as possible.

#### Participants

Fifteen healthy participants (9 females) were recruited through MRC-CBU volunteers’ panel. Their age group in this and in subsequent experiments was 18 to 40. All gave written, informed consent before the experiment, and were paid £8.50 for their participation. All had normal or corrected to normal vision.

### Results

Figure 2 and tables 1 and 2 summarize the main results. As is evident, these concur with the key predictions: trial 1 RT be the longest, trial 1 RT for 5 trial episodes > 3 trial episodes, and switch cost be absent at trial 1. The first trial of the conceived episode did take the longest to execute (table 1, and main effect of serial position in table 2). Cohen’s d (effect size: mean difference/standard deviation) was 1.68 and 1.99 for 3 and 5 trial episodes respectively. This trial 1 RT was higher for 5-trial compared to 3-trial episodes (95% CI of difference = [46, 113], table 1 and interaction between serial position and episode length in table 2, Cohen’s d = 1.3). While the performance on switch trials was poorer than on repeat trials (table 2, main effect of rule switch), this effect of rule switch differed across the serial positions within the episode (table 2: Rule Switch x Serial Position). Specifically, as predicted it was absent at trial 1 (RT: paired t_14_ = 1.05, p=0.3, 95% CI of difference = [-19, 54]; Accuracy: paired t_14_ = 1.1, p =0.3, 95% CI of difference = [-0.02, 0.01]).

**Figure 2.**
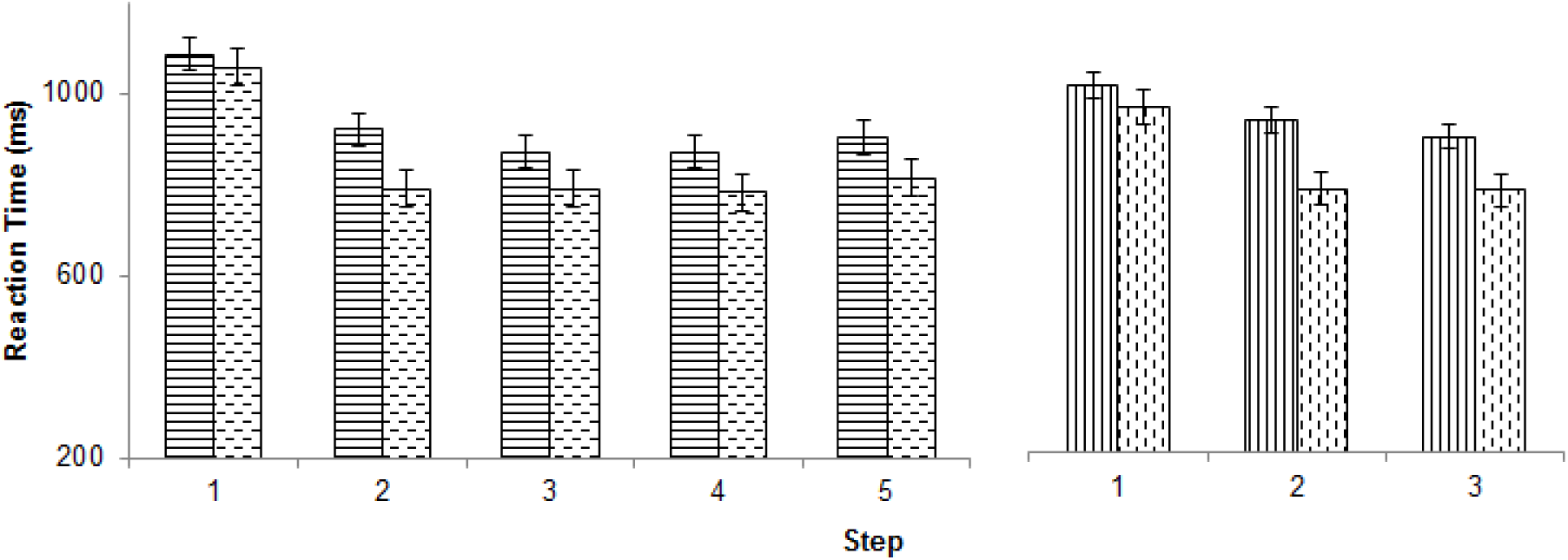
Pattern of reaction times across the steps of the 5 and 3 step episodes (continuous line bars: rule switch trials, dashed line bars: rule repeat trials). Note that trial 1 had the highest RT for both switch and repeat trials in both 5 and 3 step episodes. Error bars here and in subsequent figures represent 95% confidence intervals calculated

**Table 1.** Mean reaction times (ms) and accuracies (%, in parenthesis) across rule switch and repeat trials of 3 and 5 step episodes.

|  | 3 Step |  |  | 5 Step |  |  |  |  |
| --- | --- | --- | --- | --- | --- | --- | --- | --- |
|  | 1 | 2 | 3 | 1 | 2 | 3 | 4 | 5 |
| Switch | 1014 (93) | 938 (93) | 902 (93) | 1084 (95) | 920 (93) | 871 (94) | 870 (93) | 903 (94) |
| Repeat | 968 (94) | 785 (95) | 782 (96) | 1058 (95) | 792 (96) | 791 (95) | 783 (94) | 814 (97) |

**Table 2.**
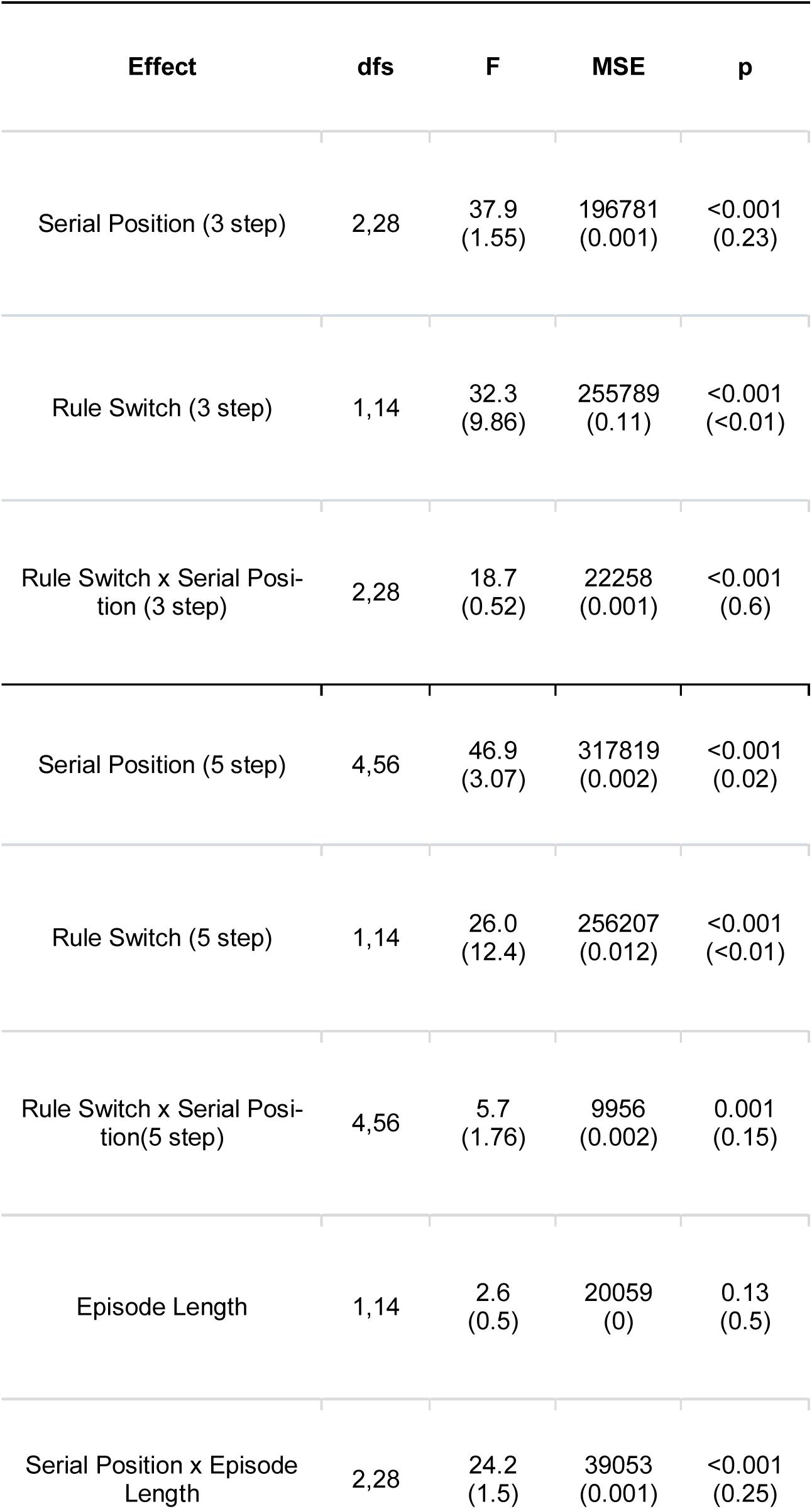
Serial Position: Repeated measures ANOVA looking at the main effect of the position of trial within the episode on RT (and accuracy, in parenthesis). Rule Switch: Main effect of rule switch (Switch vs Repeat trials). Serial Position x Rule Switch: Interaction between the effects of rule switch and serial position. Serial Position x Episode length: Effect of serial position compared across 3 step and the first three trials of 5 step episodes.

An interesting dissociation was that while elevated RTs on switch-trials were accompanied by the expected *reduced* accuracy (compared with repeat-trials), the elevated RTs on the first trial of a task episode was, if anything, associated with *greater* accuracy on those trials. This was the case even though RT on switch trials was slower only by 92 ms compared to the difference of 250 ms between trial 1 and subsequent trials. This is to be expected because trial 1 response got delayed because the episode related cognitive entity had to be assembled *prior* to the execution of trial 1, and not because of additional control demands related to the trial 1 execution.

In the next experiment, we replicated the basic findings of Experiment 1 using a different design and tested another key prediction. If trials of a task episode are executed as parts of one cognitive entity assembled at trial 1 then later trials of the episode i.e. trial 2 onwards will be executed *faster* than identical trials executed as independent tasks. This is because when trials are executed as parts of a task-episode they are partially prepared for at the beginning of the episode causing the additional delay in executing trial 1. This will cause faster execution of subsequent trials. In contrast, when trials are executed as independent tasks they will have to be individually prepared for and executed resulting in slower execution.

## Experiment 2

Here we used a different method to bias participants’ conception of what constituted a task. Series of 3 or 5 consecutive trials were grouped together into discrete task episodes by having smaller iTis (500 ms, iTi) between them, while these episodes were separated from each other by discernibly larger durations (2s; Figure 3). This was further reinforced by a faded number in the stimulus background that conveyed the number of steps left in the current episode. The first trial of a 3-trial chunk had the digit ‘3’ in the background, while the second trial had ‘2’ and the third had ‘1’. Likewise, in 5-trial chunks this background digit changed 5-4-3-2-1, across its 5 trials. Note that trials (and by extension the experiment) could be executed perfectly well without construing the series of trials as a task episode. There was nothing structural in the design that forced participants to construe the 3 (or 5) consecutive trials as parts of a higher task.

**Figure 3.**
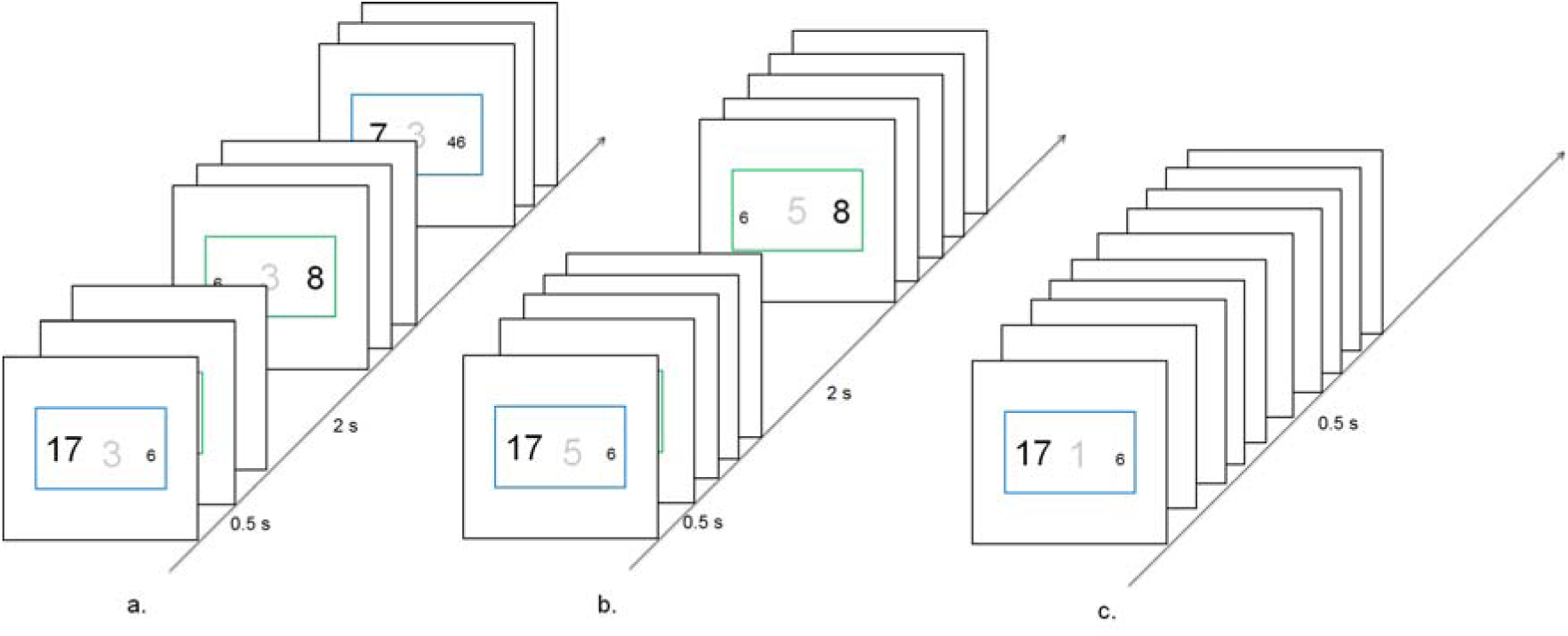
Trial rules were the same as in Experiment 1. There were three kinds of trial blocks. (a) 3 trial episodes were created by having small iTi within the episode and large iTi across episode boundaries. This was further reinforced by the faded digit in the background that went from 3-2-1 across the 3 trials making up the episode. (b) 5 trial episodes were similar to the 3 step episodes other than consisting of 5 trials. (c) Trials of the independent trial blocks were presented in a flat sequence with constant iTi and had the digit ‘1’ in their background that remained the same throughout the block.

Apart from blocks composed of 3 and 5 trial episodes there was a third block type whose trials were not organized into chunks and were instead presented as one flat sequence. Such trials had the digit ‘1’ in the stimulus background.

### Methods

Stimuli and trial rules were identical to experiment 1 (Figure 3). Before 3 and 5 step task blocks participants saw ‘3 Step Tasks’ and ‘5 Step Tasks’ respectively, onscreen till spacebar was pressed. Before the independent trial block participants saw ‘Independent Trials’, and the individual trials of this block had the number ‘1’ in the stimulus background. The 3- and 5-trial task blocks had 48 and 80 trials respectively, and the independent trial blocks had 30 trials. The order of the three block types was random. Eighteen participants (11 females) did a total of 85 – 100 blocks. They were not explicitly mentioned about the temporal grouping of the trials in the main experiment, and were asked to ignore the faded number in the stimulus background.

### Results

Figure 4, tables 3 and 4 summarize the key results. The first trial of the 3 and 5 trial episodes took longest to execute (Cohen’s d: 1.3 and 1.6; table 3, and main effect of serial position in table 4). While there was a main effect of rule switch on performance (main effect of rule switch in table 4), the effect varied across serial positions (table 4: Rule Switch x Serial Position). Specifically, as in the previous experiment it was absent on trial 1 (RT: paired t_17_ = 1.7, p=0.1, 95% CI of difference = [-8, 71]; Accuracy: paired t_17_ = 0.8, p =0.4, 95% CI of difference = [-0.01, 0.02]). Unlike the previous experiment, the trial 1 RT for 5 step episodes was not higher than 3 step episodes (95% CI of difference = [-39, 59]; table 4 interaction between serial position and episode length, Cohen’s d = 0.1). Lastly, as in the previous experiment the strong difference in RT between the first and subsequent steps was not accompanied by changes in accuracy (table 3, and main effect of serial position on accuracy in table 4).

**Figure 4.**
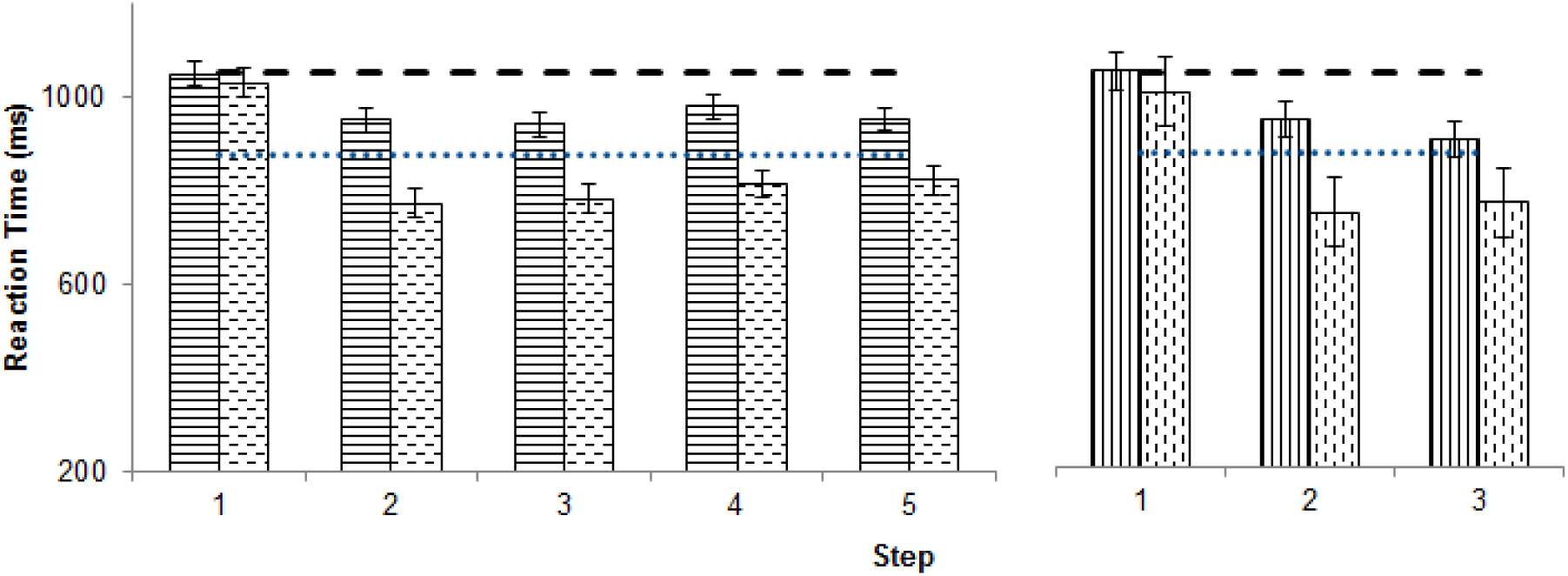
Pattern of reaction times across the steps of the 5 and 3 step episodes (continuous line bars: rule switch trials, dashed line bars: rule repeat trials). Independent dashed and dotted lines represent the switch and repeat trial RTs in the independent trial blocks. Note that these are higher than switch and repeat trial RTs of both 3 and 5 step episodes.

**Table 3.** Mean reaction times (ms) and accuracies (%) across rule switch and repeat trials of 3 and 5 step episodes, and of independent trial blocks.

|  |  | 3 Step |  |  | 5 Step |  |  |  |  | Independent trials |
| --- | --- | --- | --- | --- | --- | --- | --- | --- | --- | --- |
|  |  | 1 | 2 | 3 | 1 | 2 | 3 | 4 | 5 |  |
| RT | Switch | 1052<br>(96) | 950<br>(96) | 905<br>(94) | 1048<br>(95) | 951<br>(96) | 940<br>(94) | 979<br>(94) | 952<br>(94) | 1052<br>(95) |
|  | Repeat | 1007<br>(95) | 748<br>(95) | 769<br>(96) | 1030<br>(95) | 773<br>(97) | 782<br>(96) | 814<br>(96) | 822<br>(96) | 875<br>(98) |

**Table 4.**
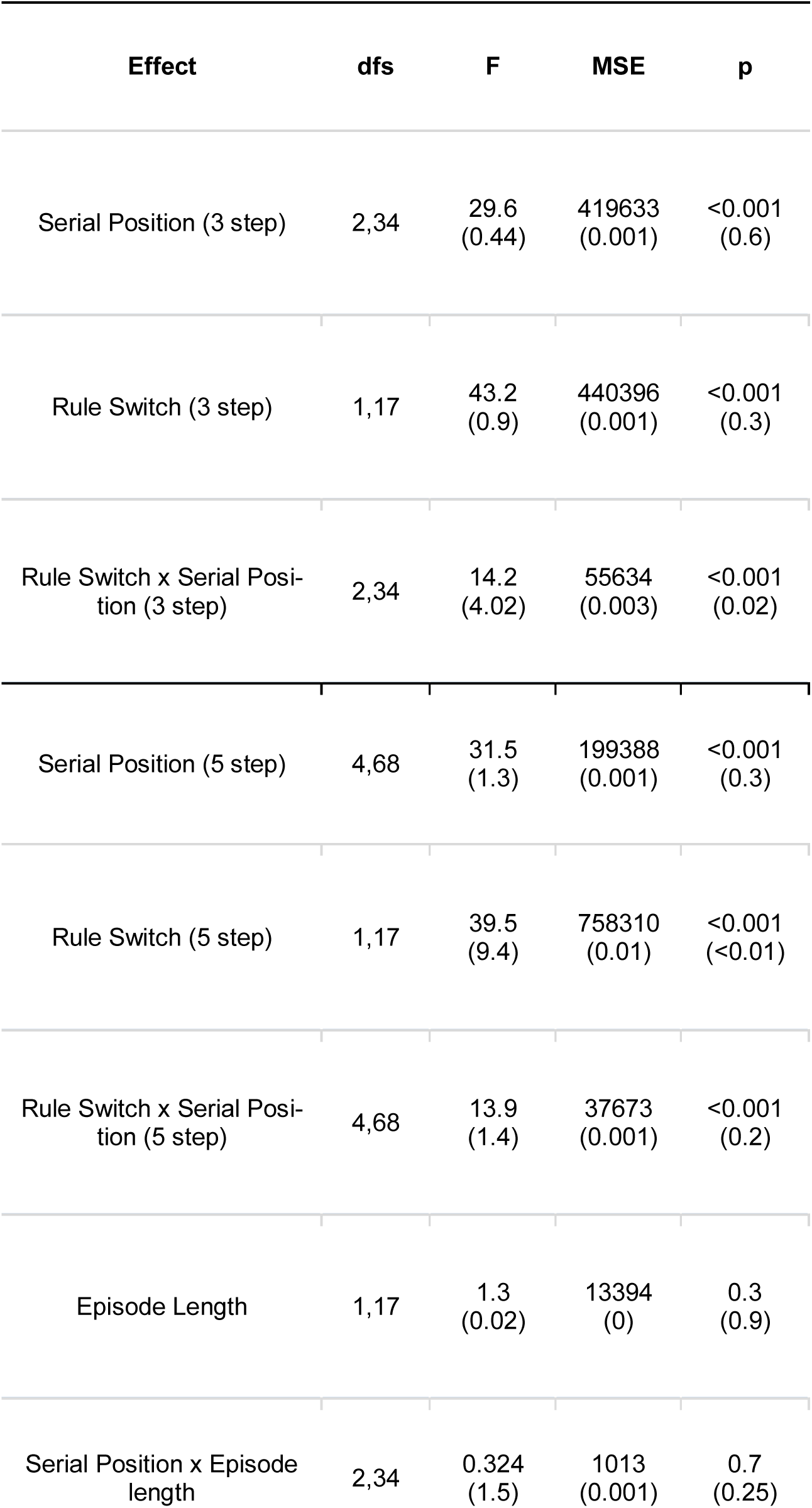
Serial Position: Repeated measures ANOVA looking at the main effect of the position of trial within the episode on RT and accuracy. Rule Switch: Main effect of rule switch (Switch vs Repeat trials). Serial Position x Rule Switch: Interaction between the effects of rule switch and serial position. Serial Position x Episode length: Effect of serial position compared across 3 step and the first three trials of 5 step episodes.

Trials conceived as parts of a task-episode were executed faster than trials conceived as independent tasks. The dashed and dotted lines in Figure 4 mark the RTs on rule switch and repeat trials in the independent trial blocks. As is evident these were higher than RTs on trials 2 and beyond of task episodes (t_17_ > 5.5, p < 0.001; Cohen’s d > 1.61). Accuracies however were not significantly different (table 3). Thus, organization of trials into larger task-episodes created a slower trial 1 but faster execution of subsequent trials. Was the price paid at trial 1 offset by gains at subsequent steps? We compared average RT and accuracy between blocks organized into task episodes with those that were not. Average RT on 3 and 5 trial task blocks was still significantly lower than on the independent trial blocks (t_17_ > 4.1, p < 0.001, Cohen’s d = 0.82). Accuracy measures, however, were not different across them. This benefit also extended to switch control. Average reaction time switch cost amongst independent trials was greater than that amongst trials executed as parts of task episodes (176 vs 128 ms; t_17_ = 2.1, p = 0.06, CI of difference = [-1, 96], Cohen’s d = 0.49).

It may be said that the above may result from participants getting more tired in independent task blocks because they were continuously executing trials while during task-episodes they got a bigger break after every few trials. This seems unlikely. Firstly, the first trial they executed after the supposed break (i.e. trial 1 of episodes) was executed much *slower* than even the independent trials. An account based on rest would have predicted that trial 1 after additional rest be executed faster than average. Further, we compared the trials of the first half of independent trial blocks with trials of the latter half of 3- and 5-trial episode blocks. Participants may be expected to be less ‘tired’ during the former (as they had executed only up to maximum of 15 trials) than latter (they would have executed at least 24 and 40 trials in 3 and 5-trial episodes respectively). This is because the independent trial blocks were significantly shorter (30 trials) compared to 3 and 5-trial episode blocks (48 and 80 trials respectively). However, results were identical to the above and participants were still faster during the task-episode trials (p < 0.01, Cohen’s d = 0.87) compared to independent trials.

Experiment 2 replicated two out of the three results of experiment 1. The reason for the absence of significantly increased RT at the beginning of 5-compared to 3-trial task episodes could have been that in the current design the same conceived task-episode was repeated across the block, hence participants did not have to organize their cognition actively enough before every episode for the small temporal difference in initiating 5 and 3 trial episodes to be evident. We tested this in experiment 3.

## Experiment 3

The design was identical to experiment 2 except that both 3 and 5 trial task episodes would now occur randomly in the same block. We expected step 1 trial differences between 3 and 5 trial task-episodes to get evident again.

### Methods

Identical to Experiment 2. The experimental session lasted half an hour during which participants did an average of 250 task episodes, which consisted of roughly equal number of 3 and 5 step episodes in a random order. 22 healthy participants (11 females) were recruited.

### Results

Tables 5 and 6 summarize the key results. Notably, the trial 1 RT was higher for 5 trial compared to 3 trial episodes (table 5 and table 6, interaction between serial position and episode length, 95% CI of difference = [15, 96], Cohen’s d = 0.6). Other results were largely a replication of the previous two experiments. The first step of the conceived episode took longest to execute (table 5, and main effect of serial position in table 6). This was the case for both switch and repeat trials at trial 1. Performance on switch trials was poorer than on repeat trials (table 6 main effect of rule switch), however, the effect of rule switch was not the same across the trials making up the episode (table 6: Rule Switch x Serial Position). Specifically, the switch cost was absent at the first position (RT: paired t21 = 0.89, p=0.4, 95% CI of difference = [-43, 109]; Accuracy: paired t21 = 0.3, p =0.7, 95% CI of difference = [-1.4, 1.9]). Lastly, accuracies were not significantly different between the trial 1 and subsequent trials (main effect of serial position on accuracy in table 6).

**Table 5.** Mean reaction times (ms) and accuracies (%) across rule switch and repeat trials of 3 and 5 step episodes, and of independent trial blocks.

|  |  | 3 Step |  |  | 5 Step |  |  |  |  |
| --- | --- | --- | --- | --- | --- | --- | --- | --- | --- |
|  |  | 1 | 2 | 3 | 1 | 2 | 3 | 4 | 5 |
| RT | Switch | 1275<br>(97) | 1125<br>(97) | 1116<br>(97) | 1333<br>(97) | 1108<br>(97) | 1112<br>(97) | 1158<br>(97) | 1154<br>(97) |
|  | Repeat | 1248<br>(96) | 925<br>(99) | 947<br>(98) | 1300<br>(96) | 904<br>(98) | 934<br>(98) | 977<br>(98) | 990<br>(97) |

**Table 6.** Serial Position: Repeated measures ANOVA looking at the main effect of the position of trial within the episode on RT and accuracy. Rule Switch: Main effect of rule switch (Switch vs Repeat trials). Serial Position x Rule Switch: Interaction between the effects of rule switch and serial position. Serial Position x Episode length: Effect of serial position compared across 3 step and the first three trials of 5 step episodes.

| Effect | dfs | F | MSE | p |
| --- | --- | --- | --- | --- |
| Serial Position (3 step) | 2,42 | 20<br>(2.9) | 797993<br>(14.1) | <0.001<br>(0.06) |
| Rule Switch (3 step) | 1,21 | 39.7<br>(2.2) | 577148<br>(27.1) | <0.001<br>(0.15) |
| Rule Switch x Serial Position<br>(3 step) | 2,42 | 11.1<br>(2.3) | 93740<br>(13) | <0.001<br>(0.11) |
| Serial Position (5 step) | 4,84 | 23.5<br>(2.5) | 697698<br>(9.9) | <0.001<br>(0.05) |
| Rule Switch (5 step) | 1.21 | 43.1<br>(6.2) | 1273614<br>(48.8) | <0.001<br>(0.02) |
| Rule Switch x Serial Position<br>(5 step) | 4,84 | 12.4<br>(1.9) | 51311<br>(7) | <0.001<br>(0.12) |
| Episode Length | 1,21 | 0.6<br>(0.6) | 5631<br>(0) | 0.5<br>(0.8) |
| Serial Position x Episode<br>length | 2,42 | 11.1<br>(0.09) | 35336<br>(0.39) | <0.001<br>(0.9) |

Current experiment affirmed that longer episodes did indeed take longer to initiate, and replicated other behavioral signatures seen in experiments 1 and 2.

## Experiment 4

Switch cost ensues from a change in implementation strategies (e.g. change in task set, recall of new rules, over-riding past task related configurations) between previous and the current acts that are conceived as different tasks (Driesbach et al. 2006; Kiesel et al. 2010; Roger & Monsell, 1995; Schneider & Logan, 2014). In experiments 1 to 3 a sequence of task items were executed as parts of a larger task episode. Cognitive accompaniments related to individual task items were subsumed by the overarching episode related cognitive entity whose dismantling/reassembly at episode boundaries left no advantage of repeating (or cost of switching) a task-item. In contrast to switch cost the congruency effect in Stroop tasks (e.g. identify the font color of the word RED when the font is blue) is not related to task identities and follows largely from the incongruence between different stimulus dimension of the current trial. It is the incongruence between the word meaning and color dimensions of the current stimulus that has to be controlled for (MacLeod, 1991; Stroop, 1935). Sequence effects on congruency effect (e.g. slower response on incongruent trials following congruent trials compared to incongruent trials following incongruent trials) are present but account for only a fraction of incongruence effects (Egner, 2007). We predicted that, in contrast to switch cost, congruence effects will not be affected by the dynamics of task-episode related entity and hence will *not* be absent at trial 1 of task-episodes.

Participants did a modification of color-word Stroop task (Figure 5). The experiment had two kinds of blocks. Trials of the first one were temporally grouped into task episodes similar to that in Experiments 2 and 3. In such blocks four consecutive trials were grouped together through smaller iTi (500 ms), while two adjacent episodes were separated by longer intervals (1500 ms). Furthermore, the stimulus background had a faded digit that moved 4-3-2-1 across the four trials of the episode.

**Figure 5.**
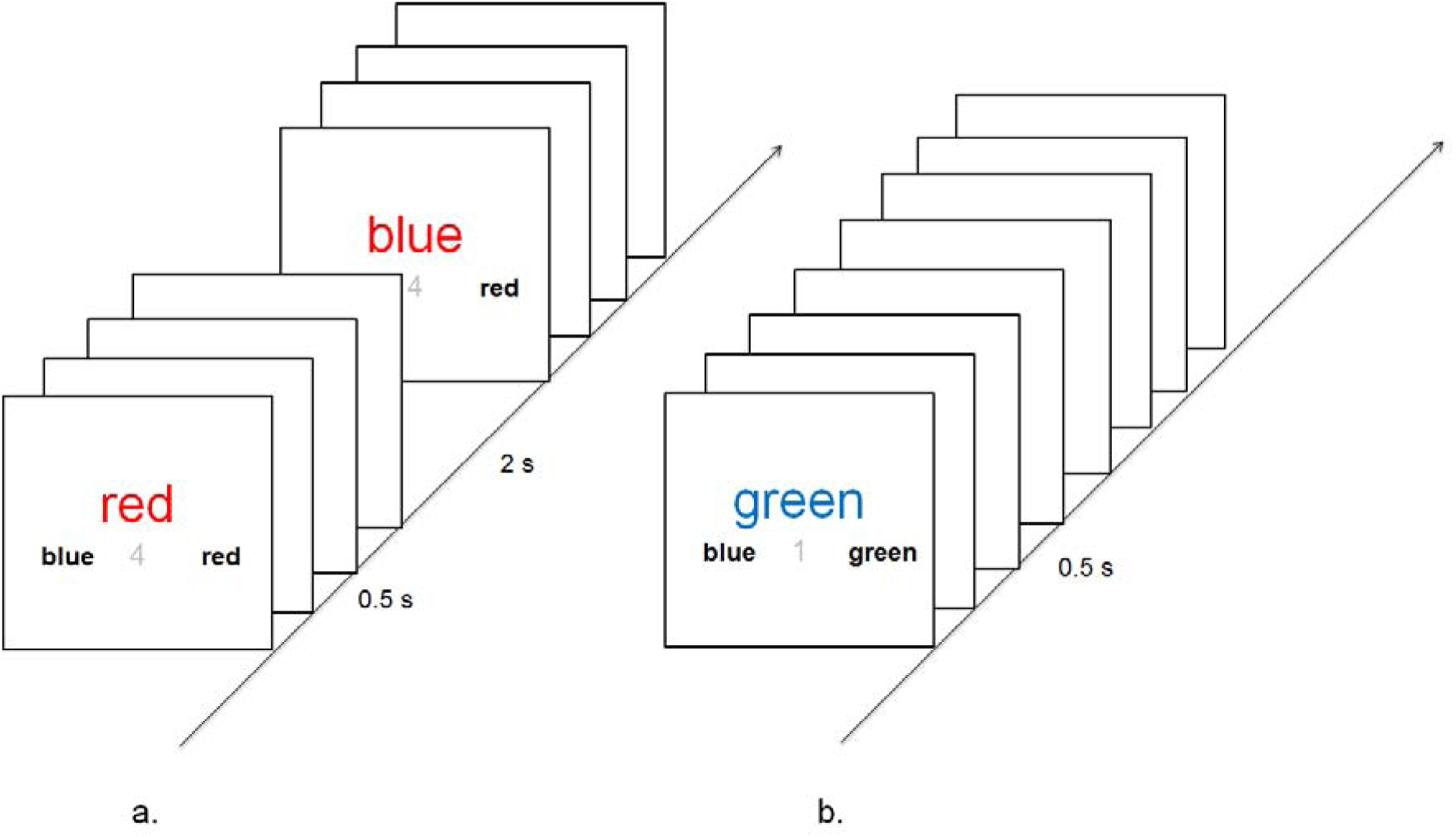
Participants chose the color of the print of the central word from the two choices below. Notion of task episodes was created by having short iTi between the four trials making up the supposed episode and long iTi between the consecutive trials across successive episodes. Additionally, a faded number went from 4 to 1 across the four trials of these episodes. In a second block type trials were presented as a flat sequence with constant iTi across the block. All trials of such blocks had the number ‘1’ in their background.

In the second kind of blocks, trials were not organized into larger task episodes and were presented as a flat sequence (iti = 500 ms) with the digit ‘1’ in the stimulus background. These blocks allowed us to replicate the results of experiment 2 that had shown faster execution of trials executed as parts of a larger task episode than independent trials.

### Methods

On each trial participants saw a centrally presented color name (‘Red’, ‘Blue’, ‘Green’, ‘White’; in Arial font size 40) in colored prints, such that the print and the name were congruent on 60% of trials and incongruent on the rest (Figure 5). Participants chose the color of the print from the two color words presented in Arial black font size 20 (1 degree away from the center and ½ degree below it) below by pressing a button spatially congruent with their choice - left: Numpad 1 (to be pressed with right index finger); right: Numpad 2 (to be pressed with right middle finger). One (allocated to left or right at random) always reflected the font color. When the font and word were incongruous, the other word always repeated the stimulus word. On congruous trials the other word was selected randomly from the remaining colors. Between these two options a partially (70%) transparent digit appeared in black. The stimuli remained on screen till a response was made. Erroneous responses elicited a low-pitched feedback tone.

At the start of 4-trial blocks the instruction screen mentioned ‘4 - step blocks’, it remained on until the spacebar was pressed. Such blocks consisted of 48 trials. The instruction for the independent trial block mentioned ‘independent trials’. Such blocks consisted of 20 trials. These two blocks were interleaved. Eighteen participants (9 females) did a total of 150 - 250 blocks.

### Results

Figure 6 and tables 7 and 8 summarize the key results. As before, the first trial of the episode took longest to execute (table 7, and main effect of serial position in table 8, 95% CI of difference = [54, 146], Cohen’s d = 1.1). This was the case for both congruent and incongruent trial 1. Expectedly, the performance on incongruent trials was poorer than on congruent trials (table 8 main effect of rule switch), but crucially it did not vary across the episode (table 8: Rule Switch x Serial Position). Specifically, unlike switch cost in previous experiments, the cost of incongruence did not disappear on trial 1 of the episode (RT: one-sample t_17_ = 8.8, p=<0.001, 95% CI of difference = [127, 206]; Accuracy: one-sample t_17_ = 4.2, p = <0.001, 95% CI of difference = [1.7, 5.0]).

**Figure 6.**
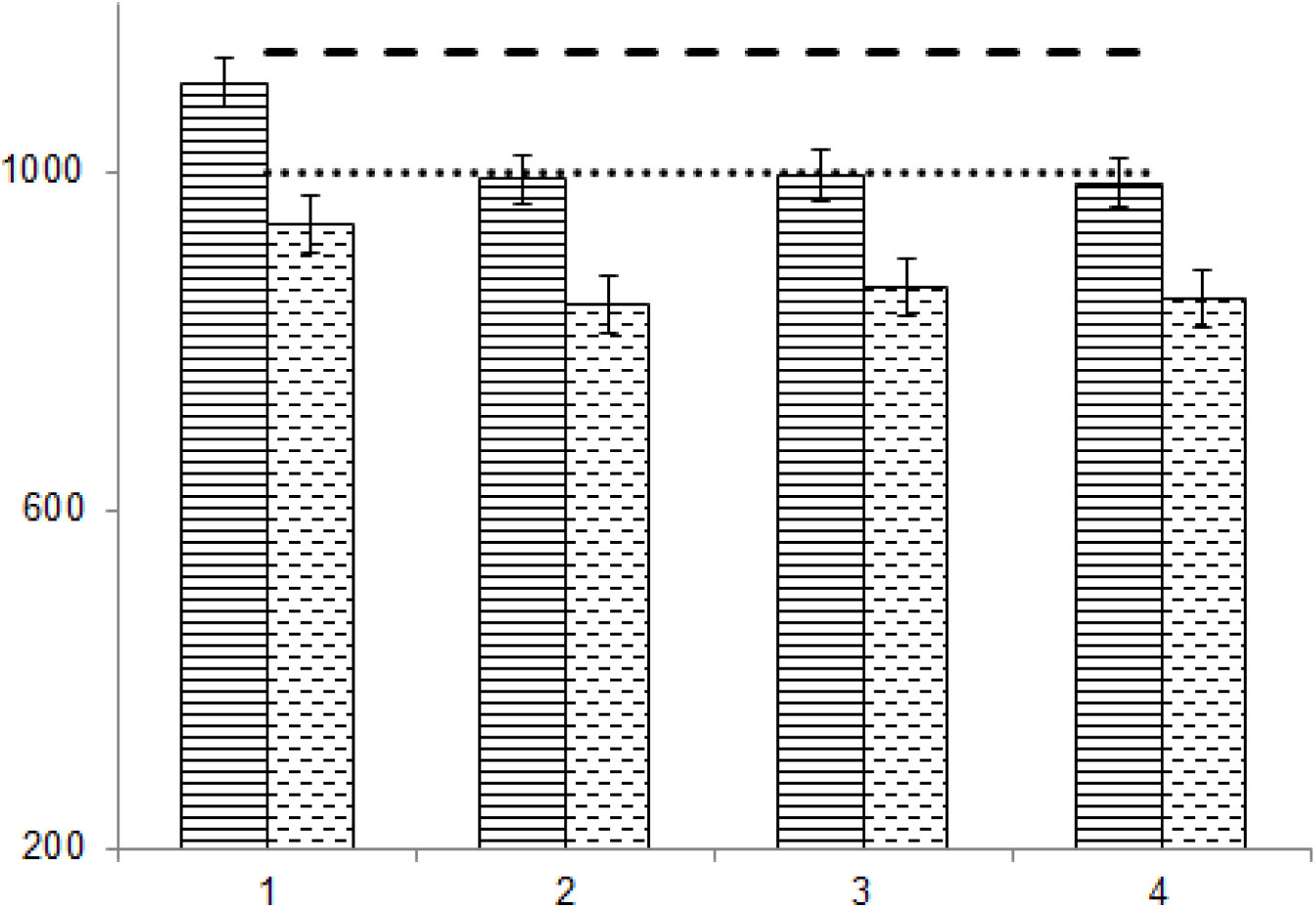
Pattern of reaction times across the four trials of the construed task episodes (continuous line bars: incongruent trials, dashed line bars: congruent trials). Independent dashed and dotted lines represent the incongruent and congruent trial RTs in the independent trial blocks. Note that these are higher than the corresponding trials that formed part of a larger task episode.

**Table 7.** Mean reaction times (ms) and accuracies (%) across the incongruent and congruent trials corresponding to the four steps of task episode and of independent trial blocks.

|  |  | 1 | 2 | 3 | 4 | Independent tri-<br>als |
| --- | --- | --- | --- | --- | --- | --- |
| RT | Incongru-<br>ent | 1104<br>(96) | 991<br>(94) | 995<br>(94) | 986<br>(94) | 1141<br>(92) |
|  | Congruent | 938<br>(99) | 843<br>(99) | 862<br>(99) | 848<br>(99) | 1000<br>(99) |

**Table 8.**
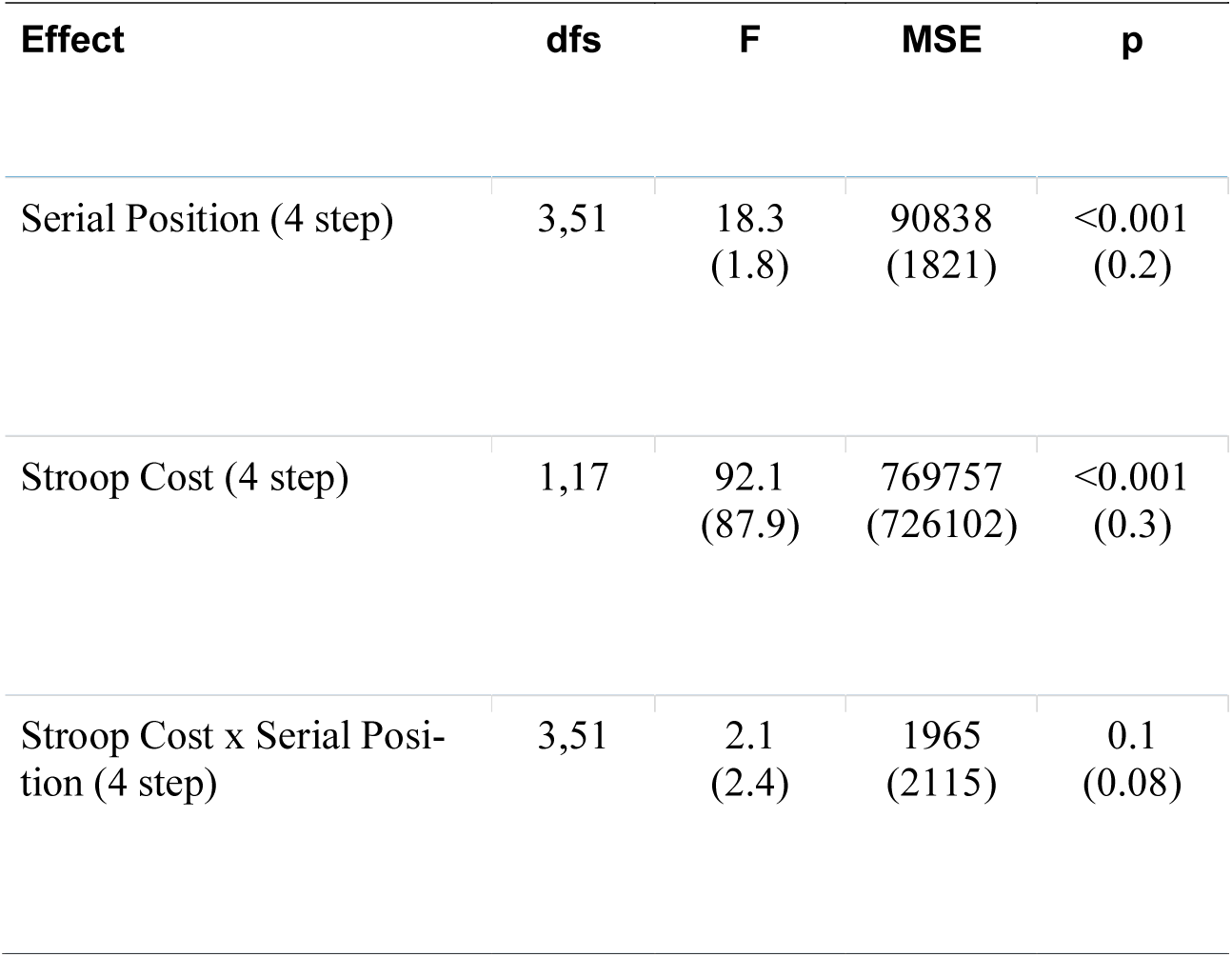
Serial Position: Repeated measures ANOVA looking at the main effect of the position of trial within the episode on RT (and accuracy, in parenthesis). Stroop Cost: Main effect of Congruence (Incongruence vs Congruence trials). Stroop Cost x Serial Position: Interaction between the effects of congruence and serial position.

| Effect | dfs | F | MSE | p |
| --- | --- | --- | --- | --- |
| Serial Position (4 step) | 3,51 | 18.3<br>(1.8) | 90838<br>(1821) | <0.001<br>(0.2) |
| Stroop Cost (4 step) | 1,17 | 92.1<br>(87.9) | 769757<br>(726102) | <0.001<br>(0.3) |
| Stroop Cost x Serial Position (4 step) | 3,51 | 2.1<br>(2.4) | 1965<br>(2115) | 0.1<br>(0.08) |

The dashed and dotted lines in Figure 6 mark the RTs on incongruent and congruent trials in the independent trial blocks. As is evident both were higher than the incongruent and congruent trials of task episodes. Infact the average RT on trials executed as parts of task episodes was lower than for independent trials even when task episode trial 1 RTs were included in comparison (congruent trials: t_17_ = 7, p<0.001, 95% CI of difference = [89, 165]; incongruent trials: t_17_ = 4.9, p<0.001, 95% CI of difference = [69, 175]). Whilst there was a general RT advantage for task episode trials, Stroop cost on RT (RT on incongruent trials - RT on congruent trials) did not differ between task episode and independent trials (t_17_ = 0.3, p=0.8, 95% CI of difference = [-27, 37]). However, Stroop cost on accuracy (Accuracy on Congruent trials - Accuracy on incongruent trials) was significantly higher on independent than task episode trials (t_17_ = 2.6, p=0.02, 95% CI of difference = [0.5, 5.5]). As might be expected from that result, when Stroop trials were broken down by congruency, accuracy was greater for task episode than individual trials for incongruent trials (t_17_ = 2.7, p=0.02, 95% CI of difference = [0.6, 5.3]), but there was no difference on congruent trials (t_17_ = 0.07, p=0.9, 95% CI of difference = [-0.5, 0.5]).

These results confirmed the prediction that Stroop cost would not be absent on trial 1, and that trials executed as parts of a task episode were executed faster and better controlled than trials executed as independent tasks.

## Experiment 5

Counting or countdowns involved in prior experiments can be construed as a hierarchical act, and it is possible that the hierarchy evident in the execution of trials in experiments 1 to 4 was a spillover from a separate but simultaneous hierarchical task – counting (or countdown) in 3s, 5s or 4s. Through this experiment we clarify that this was not the case. Instead the above results were the consequence of the construal of the trial series as one task entity.

The task episodes of the current experiment did not have a fixed number of trials and did not involve counting or countdown. The idea of episodes was created by the presence of an additional margin (outside the one conveying the relevant trial rule) that stayed on for an extended duration during which trials would appear randomly at any time. Trials would not appear after this margin had switched off. Every episode began with the appearance of trial 1 stimulus along with this margin (Figure 7). While the trial 1 stimuli disappeared after the response, the outer margin remained on. Subsequent trials would appear anytime while this outer margin was on. This margin went off marking the end of the episode, and came back on with the beginning of the next episode. The notion of task episode was thus implicitly defined as the period enveloped by the presence of the outer margin.

**Figure 7.**
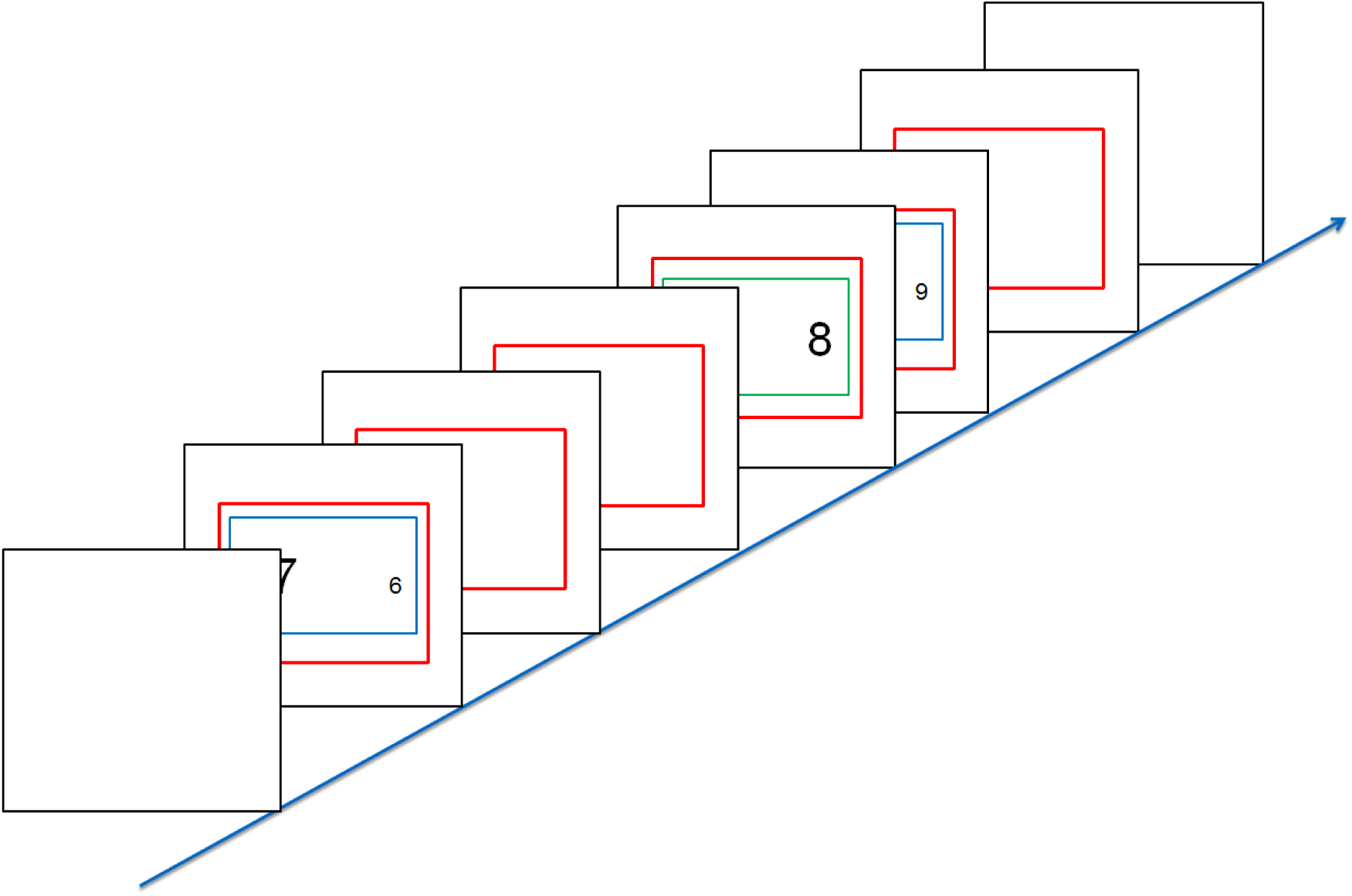
The outermost margin (here in red) stayed on for extended duration during which an unpredictable number of trials would appear at random intervals. Trials would not appear when this margin was off. We hoped that participants would construe the temporal epochs carved by the presence of such margins as the recurring task episodic units to be executed.

Two kinds of episodes (short and long), framed by different color margins (black or red), were randomly interleaved. Short episodes lasted between 3 to 6 seconds and two to three trials would appear at any time during this period. Long episodes lasted between 7 to 10 seconds and three to seven trials could appear during this period. The inter-episode interval was fixed at 2 s, but the iTi within an episode varied from 50 ms to 5 s in short episodes and 50 ms to 3 s in long episodes. Note that in this experiment, not only did the subjects not have any foreknowledge about the identity and sequence of task items, they also could not predict the number of trials and their iTi.

### Methods

Apart from the additional outer margin, individual trials were identical to Experiments 1 - 3 (speeded decision as to which of two numbers had the smallest value or font size as indicated by the color of the inner margin). 27 participants (17 females) did two experimental sessions each lasting 16 minutes, with a period of rest in between. Within a session the two kinds of conceived episodes occurred randomly. Participants were not instructed about the episodic structure of the task block and were told nothing about the outermost margins.

### Results

Key results of previous experiments were again replicated (Table 9 and 10). The first trial of the episode took longest to execute (F_2, 52_ = 28.1, p < 0.001). This trial 1 RT was longer before longer episodes (95% CI of difference = [14, 39], Cohen’s d = 0.81; t_26_ = 4.2, p < 0.001). Switch cost was substantially reduced at the first trial of the episode (Smaller episodes: F_2, 52_ = 18; Longer episodes: F_6, 156_ = 8.1, p < 0.001 for both). The execution of trials construed as parts of a larger task episode again showed key behavioral signatures that suggested that these trials were executed not as independent entities but as parts of a routine. This was the case even though participants were unaware of the number and sequence of task items they would be required to perform, when those items would be executed and the precise duration of the episode.

**Table 9.** Mean reaction times (ms) and accuracies (%) across the switch and repeat trials of short and long episodes.

|  |  | Short |  |  | Long |  |  |  |  |  |  |
| --- | --- | --- | --- | --- | --- | --- | --- | --- | --- | --- | --- |
|  |  | 1 | 2 | 3 | 1 | 2 | 3 | 4 | 5 | 6 | 7 |
| RT | Switch | 873<br>(94) | 876<br>(94) | 876<br>(92) | 898<br>(92) | 853<br>(92) | 883<br>(94) | 867<br>(94) | 874<br>(92) | 851<br>(93) | 886<br>(92) |
|  | Repeat | 843<br>(95) | 723<br>(97) | 751<br>(97) | 856<br>(94) | 723<br>(96) | 722<br>(97) | 741<br>(95) | 755<br>(95) | 775<br>(95) | 746<br>(97) |

**Table 10.** Serial Position: Repeated measures ANOVA looking at the main effect of the position of trial within the episode on RT (and accuracy, in parenthesis). Rule Switch: Main effect of rule switch (Switch vs Repeat trials). Serial Position x Rule Switch: Interaction between the effects of rule switch and serial position. Serial Position x Episode length: Effect of serial position compared across short and the first three trials of long episodes.

| Effect | dfs | F | MSE | p |
| --- | --- | --- | --- | --- |
| Serial Position (short) | 2,52 | 28<br>(1.2) | 163851<br>(0.002) | <0.001<br>(0.3) |
| Rule Switch (short) | 1,26 | 119<br>(17.9) | 544179<br>(0.05) | <0.001<br>(<0.001) |
| Rule Switch x Serial Position (short) | 2,52 | 18<br>(4.7) | 51458<br>(0.007) | <0.001<br>(0.01) |
| Serial Position (long) | 6,156 | 11<br>(2) | 38664<br>(0.004) | <0.001<br>(0.07) |
| Rule Switch (long) | 1,26 | 135<br>(16.2) | 1243930<br>(0.09) | <0.001<br>(<0.001) |
| Rule Switch x Serial Position (long) | 6,156 | 8.1<br>(1.4) | 19205<br>(0.002) | <0.001<br>(0.2) |
| Episode Length | 1,26 | 0.02<br>(1.3) | 87<br>(0.003) | 0.9<br>(0.3) |
| Serial Position x Episode Length | 2,52 | 6.1<br>(0.07) | 3840<br>(0.4) | <0.01<br>(0.9) |

## Experiment 6

In this experiment, we investigated two questions. First, is the higher trial 1 RT during 5 - trial episodes compared to 3-trial episodes caused by greater number of trials or the greater duration of the episode? Second, is the difference in RTs between longer and shorter episodes limited to trial 1? Through these we investigated if the magnitude of the subsuming episode related entity is related to the temporal length of the episode, and whether maintaining this entity impinges upon the limited cognitive reserves available for trial execution.

The first question could not be answered by previous experiments because episodes with greater number of trials were also greater in temporal duration. Regarding the second question experiments 1 to 5 gave a null result. While trial 1 RT was longer before longer episodes, RTs on subsequent trials were not different across longer and shorter episodes (effect of episode length vs serial position x episode length in Tables 2, 4, 6, 10). Thus, the maintenance of the larger episode related entity did not seem to require additional limited capacity reserves that would have decreased their availability for trial execution. It is possible that this absence of evidence was the result of *passive* nature of episodes in these experiments. Participants were merely biased to construe the set of trials as an episode. The difference in the magnitude of the subsuming episode related entity between 5 and 3 trial episodes may not have been large enough to cause discernible depletion of cognitive reserves available for component trial execution.

In the current experiment participants had to *actively* execute the task episode. Participants now had to keep an active count of trials they executed and press a ‘task end’ button at the end of each episode. Participants executed trials that were identical to those in experiments 1-3 in three different kinds of blocks - 3-trial-short, 5-trial-short and 3-trial-long (Figure 9). In the first two there was no iTi within an episode and the next stimulus onset immediately after response, whereas in the third (i.e. the 3-trial-long episodes) the iTi was 2 s.

**Figure 9:**
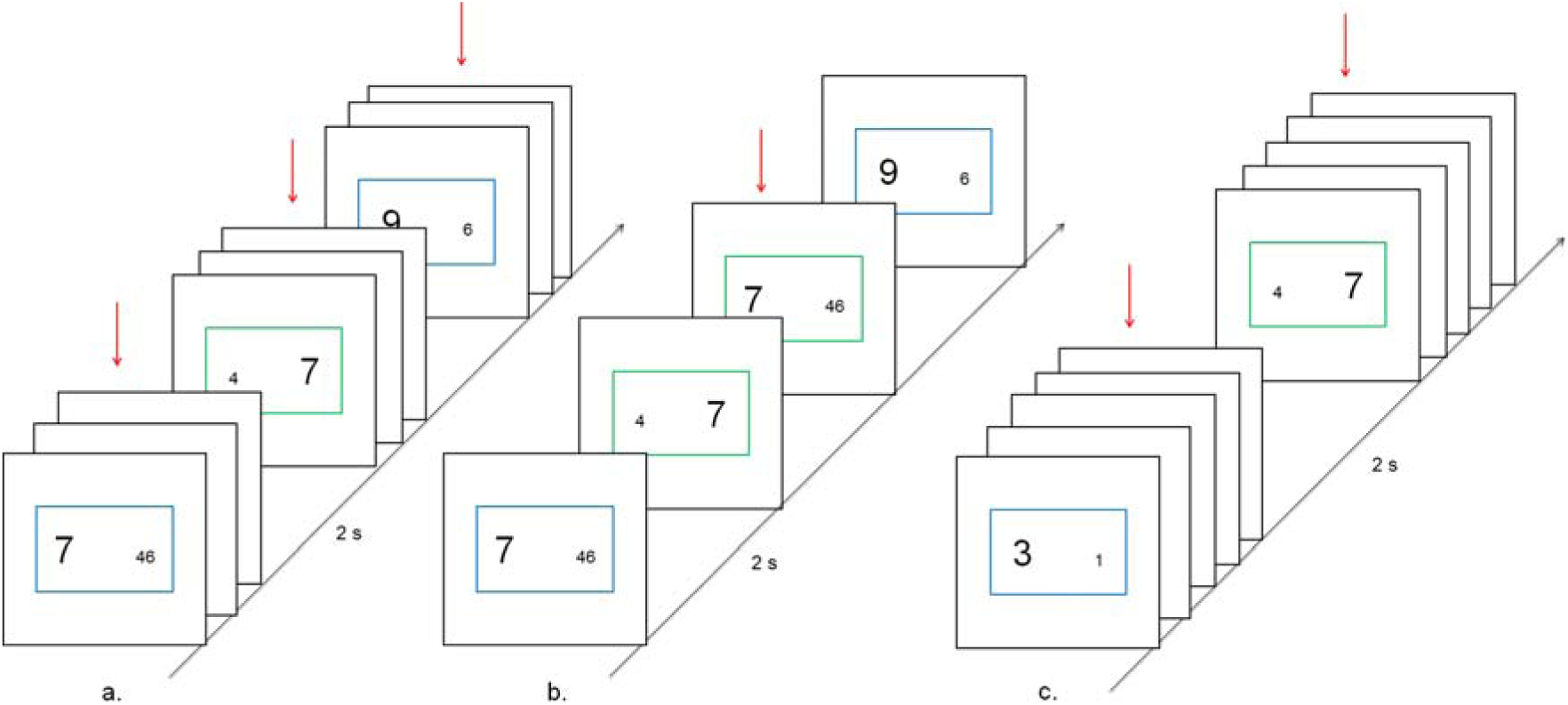
There were three kinds of blocks 3-trial-short, 5-trial-short and 3-trial-long. iTi within an episode in the first two were 0 s and in the third 2 s. Inter-episode interval was 2 s in all conditions. To ensure that the trials were actively executed as one episode participants had to keep a count of trials they had executed and then press an additional ‘task end’ key at the completion of each episode.

We made two predictions. First, if the episode related entity subsumes cognitive processing during the entire period of the task episode then its magnitude may be related to the duration of the episode and not just the number of component trials. As a result, RT on trial 1 of 3-trial-long episodes may be slower than those of 3-trial-short episodes. Second, if maintaining the episode related entity required limited cognitive resources then more of these resources will be used up during longer episodes. Less of these will be available for executing individual trials. RTs on trial 2 onwards will be slower on longer compared to shorter episodes.

### Methods

Stimuli and trial rules were identical to Experiment 1. The key differences in this experiment concerned iTi and the additional requirement to press a ‘task end’ button (key ‘Z’ on QWERTY keyboard) at the completion of each task episode. In 3-trial-short and 5-trial-short episodes, the stimuli for the next trial onset immediately after the response to the previous trial (Figure 9a and b). Within 3-step-long episodes the onset of the next trial occurred 2 s after the response to the previous trial during which the monitor was blank (Figure 9c). In all conditions there was a 2 s delay between the end of an episode and the beginning of the next. In the experiment session, blocks comprised 60 trials (i.e. 20 task episodes in 3-trial-short and 3-trial-long blocks and 12 task episodes in 5-trial-short blocks). Nineteen Participants (11 females) completed 30-40 blocks, the different number reflecting how long they chose to rest between in between blocks in the fixed-duration experimental session. Block order was randomized for each participant.

### Results

Behavior during 3-trial-short and 5-trial-short episodes was largely identical to that seen in previous experiments (Tables 11 and 12). The key difference was in their comparison. Recall that in previous experiments only trial 1 RT during 5-trial episodes was longer than 3-trial episodes.

**Table 11.**
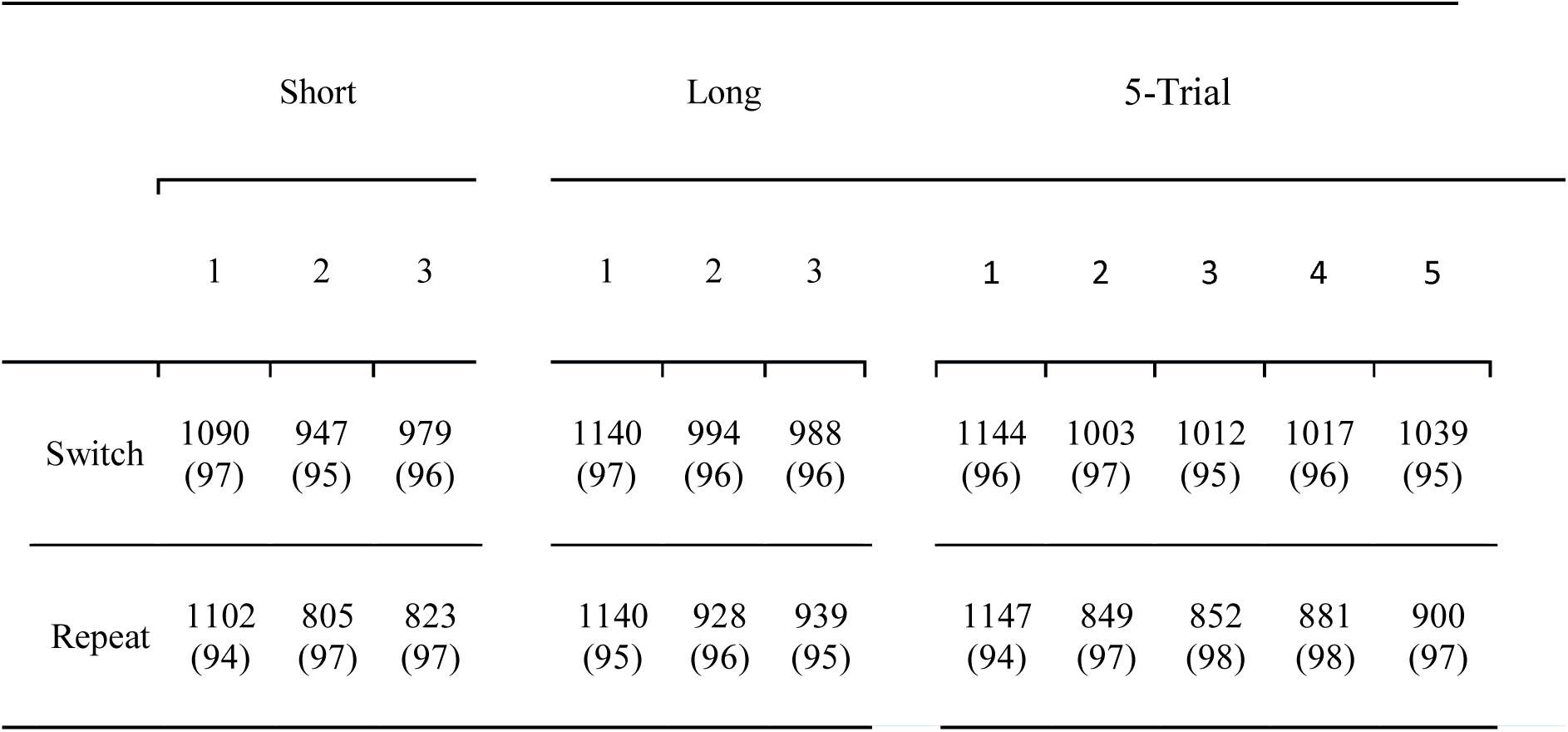
Mean reaction times (ms) and accuracies (%) across the switch and repeat trials of the three task episodes.

**Table 12.**
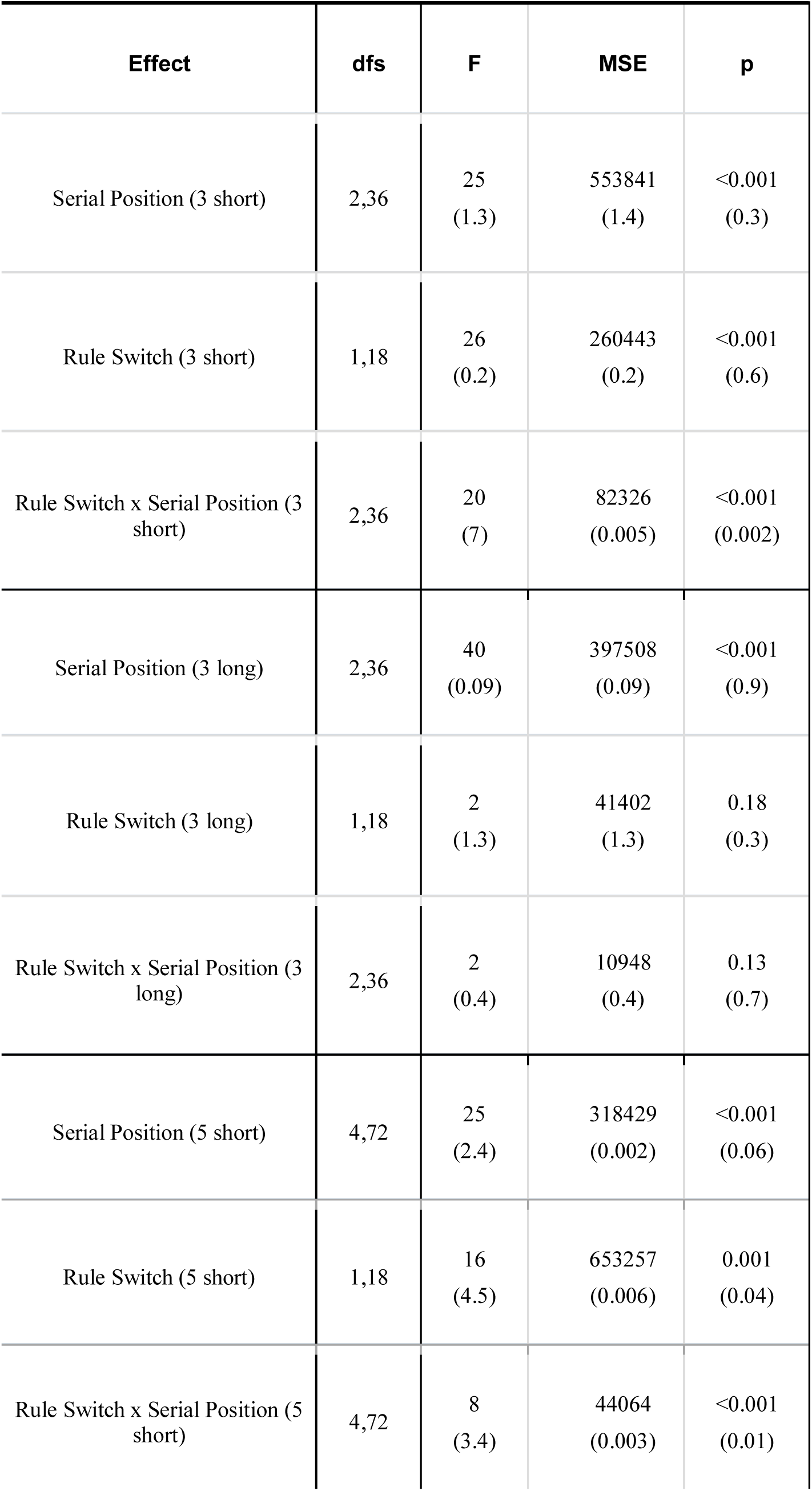
Serial Position: Repeated measures ANOVA looking at the main effect of the position of trial within the episode on RT (and accuracy in parenthesis). Rule Switch: Main effect of rule switch (Switch vs Repeat trials). Serial Position x Rule Switch: Interaction between the effects of rule switch and serial position.

| Effect | dfs | F | MSE | p |
| --- | --- | --- | --- | --- |
| Serial Position (3 short) | 2,36 | 25<br>(1.3) | 553841<br>(1.4) | <0.001<br>(0.3) |
| Rule Switch (3 short) | 1,18 | 26<br>(0.2) | 260443<br>(0.2) | <0.001<br>(0.6) |
| Rule Switch x Serial Position (3 short) | 2,36 | 20<br>(7) | 82326<br>(0.005) | <0.001<br>(0.002) |
| Serial Position (3 long) | 2,36 | 40<br>(0.09) | 397508<br>(0.09) | <0.001<br>(0.9) |
| Rule Switch (3 long) | 1,18 | 2<br>(1.3) | 41402<br>(1.3) | 0.18<br>(0.3) |
| Rule Switch x Serial Position (3 long) | 2,36 | 2<br>(0.4) | 10948<br>(0.4) | 0.13<br>(0.7) |
| Serial Position (5 short) | 4,72 | 25<br>(2.4) | 318429<br>(0.002) | <0.001<br>(0.06) |
| Rule Switch (5 short) | 1,18 | 16<br>(4.5) | 653257<br>(0.006) | 0.001<br>(0.04) |
| Rule Switch x Serial Position (5 short) | 4,72 | 8<br>(3.4) | 44064<br>(0.003) | <0.001<br>(0.01) |

Hence, only the interaction between effects of serial position and episode length was significant, and the main effect of episode type on RT was not significant. But, in the current experiment the main effect of episode was significant (F_1, 18_ = 10.5, p =0.004) but its interaction with serial position was not (F_2, 36_ = 0.4, p = 0.6). Thus, not only trial 1 but all RTs on all trials of 5-trial-short episodes were slower than 3-trial-short episodes.

Same was found for the comparison between 3-trial-short and 3-trial-long episodes (F_1, 18_ = 13, p =0.002). However, this comparison carries the caveat that trials 2 and 3 of 3-trial-long episodes were preceded by very different iTis. The effect of iTi duration on RT in the context of episodes is not known and hence the slower trials 2 and 3 RTs in 3-trial-long episodes cannot be exclusively attributed to the greater load of their episode related cognitive entity. Nonetheless, trial 1 of these episodes were matched in their preceding iTis and hence any difference between them could only be attributed to the episode they began. That this was longer for the 3-trial-long episode (paired t_18_ = 2.9, p= <0.01, 95% CI of difference = [17, 103], Cohen’s d = 0.67) suggested that the magnitude of the episode related entity was affected not just by the number of constituent trials or steps but also by the temporal duration of the episode.

The results made two suggestions. First, that at least in some conditions maintaining the episode related cognitive entity may decrease limited capacity cognitive reserves available for the execution of component step. Second, that the magnitude of this entity is related not only to the number of steps in the episodes but also to their total temporal duration.

## Discussion

We showed that every time a series of otherwise independent and unpredictable trials was construed as a task episode (1) trial 1 had the longest RT, (2) which was longer before longer episodes, and (3) trial item related switch cost was specifically absent when the switch crossed construed episode boundaries. Results (1) and (2) showed that executing trial 1 required additional processing that was related not to the intrinsic demands of trial 1 but to the episode begun by this trial, and suggested that some episode-related cognitive entity had to be assembled every time a task-episode was to be executed. (3) suggested that this episode related cognitive entity had a hierarchical relation with cognitive configurations related to component trials, such that changing the episode related entity necessarily changed or ‘refreshed’ the trial related configurations. As a result repeating and switching a trial type across episode boundaries became identical i.e. in both cases trial related configurations (e.g. trial related set) had to made afresh.

These behavioral evidence of assembly of episode-related entities at episode beginning are complemented by the neuroimaging evidence of their dismantling at episode completion. Completion of action sequences, perceptual episodes, task episodes typically elicit additional activity (Farooqui et al. 2012; Fuji & Graybiel, 2003; Zacks et al. 2001; see also Fox et al. 2005; Konishi, Donaldson & Buckner, 2000). This additional activity can again not be attributed to the intrinsic characteristics of the last event but only to the episode being completed. Hence, for example, its magnitude and spread is affected by the hierarchical level of the episode completed (subtask < task).

fMRI of experiments 2 and 3 (Farooqui et al. under review) showed that in neural regions known to deactivate in response to cognitive load (e.g. the Default Mode Network, Fox et al. 2005) beginning a task episode elicited a strong deactivation that then gradually decreased and activity returned to baseline as the sequential steps of the episode were executed. This sequential *decrease* in *deactivation* across sequential steps of the episode suggested that some cognitive entity changed as parts of the episode were executed. Again, this entity had to be related to the episode and not to the constituent trials because these trials were identical to each other. Plausibly, this sequential change in activity was related to the gradual dismantling of the episode related entity as related parts of the episode were executed.

It is also evident that this cognitive entity was not limited to propositional symbolic task sequence representation. Such a representation was absent. It may be claimed that the knowledge that the episode consists of 3 or 5 trials was the task sequence representation. This however does not explain why trial 1 RT remained high even when the same episode was iteratively executed? It is unlikely that this knowledge and its corresponding representation was forgotten at the end of every episode and had to be recalled afresh for the next iteration. This view can also not explain why recalling that the episode will be long or consist of 5 trials take longer than recalling that episode will be short/consist of 3 trials. Finally, this can also not explain why beginning a long 3-step episode should take longer than beginning identical but shorter 3-step episode.

In summary, taken together, the current and related studies (Farooqui et al. 2012; Farooqui et al. under review) suggested that goal-directed behavior is subsumed by cognitive entities related to the conceived task and goal identity through which behavior is being executed. These entities are assembled at the beginning of task episode execution, manifesting in delayed trial 1 RTs. They contain elements related to the entire duration of the episode, causing longer trial 1 RTs during longer compared to shorter episodes. These dismantle as related parts of the episode are executed causing gradual change in related activity across the duration of the task episode (Farooqui et al. under review). Completion of the episode dismantles them completely and elicits widespread activity (Farooqui et al. 2012). Insights into the nature of this entity will have to await further studies, what follow are our current speculations.

### Programs

A consequence of the hierarchical nature of our actions is that we execute extended sequence of cognition and behavior as one entity, e.g. the various components of checking email at different levels of detail (‘open browser’, ‘move cursor’, ‘click’) are executed as parts of one task and not as independent acts (e.g. Botvinick, Niv & Barto 2009; Dezfouli, Lingawi, & Balleine, 2014; Logan, 1988). An extended episode of behavior or cognition can be executed as one only through the intermediation of some subsuming entity that is selected and instantiated as a unit but corresponds to or controls over an episode of behavior and cognition. Accounts of cognition have proposed constructs like plans, programs, scripts, schema and frames that subsume and control extended behavior (Cooper & Shallice, 2000; Miller et al. 1960; Minsky, 1975; Schank & Abelson, 1977). However, these accounts have typically emphasized the hierarchical nature of the representational content of these constructs that corresponds to an abstract knowledge of what to do when in the task situation. The contained sequence of representations in these constructs is proposed to control the identity and sequence of component steps. A script about visiting a restaurant may consist of ‘find a seat’, ‘read the menu’, ‘order food’; each of which may have their own subscripts (Schank & Abelson, 1977). A schema for sweetening a cup of coffee may consist of picking up a spoon, dipping the spoon into some sugar contained within a sugar-bowl, filling the spoon with sugar, transporting it and then tipping it into the mug (Shallice & Cooper, 2000). In comparison, the current results evidenced hierarchical cognitive entities in situations where knowledge of what to do when was not available and task knowledge was not organized into hierarchical representations.

We think that the episode related entities evidenced here are the cognitive means of hierarchical execution and may be seen as a *program* through which cognition automatized parts of task episode execution that can be automatized. For example, when an extended task is intended changes related to goal, task focus and attention to be made across time in the various domains of cognition and behavior can be made in one-go by instantiating such a program that will then automatically bring about such changes across the length of the task episode. This may explain not only why trial 1 of the episode took longer to execute and that this trial 1 RT was related to the expected length of the incoming task episode. It may also explain why switch cost was absent at trial 1. Because trials of the episode were executed as parts of a program, benefits of repeating an item would be apparent within a program and not across programs. The current study beyond evidencing the presence of such programs, does not tell much about their nature and content, which will have to await subsequent studies. Following are some of the possibilities that are not mutually exclusive.

Possibility 1: The entity was a means of *organizing* cognition into task focused cognitive episodes. Through it various irrelevant processes and representations were cleared out from a position where they could compete for cognitive reserves or disrupt task execution, and were maintained in abeyance across time. Various task relevant learnings, memories, skills, dispositions, knowledge and expectancies, and the corresponding configurational changes in various perceptual, attentional, mnemonic, and motor processes may be brought to fore (e.g. Bartlett, 1932; Dashiel, 1940; Logan & Gordon, 2001; Mayr & Keele, 2000; Miller, Galanter & Pribram, 1960; Rogers & Monsell, 1995). The episode related entity embodied the program to achieve these widespread changes across time. E.g. in current task episodes processing related to mind-wandering, other ongoing unconscious goals, task irrelevant sensory and motor processing etc. had to be relegated. At the same time, the predictiveness of the episode was to be utilized to make anticipatory changes; e.g. knowledge that responses were right handed, visual attention limited to area around fixation, along with an implicit idea of iTis, could be used increase there preparativeness at times when a stimulus was expected and decrease when iTi was expected.

Possibility 2: This entity embodied the goal. Rather than seeing goal as merely an end state to be achieved, goals may be better conceived as cognitive structures or programs geared towards reaching that end state. Indeed, a wide variety of frameworks do accept that goals are important for the control and execution of tasks that lead to their achievement (Anderson, 2014; Greenwald, 1972; Gollwitzer & Sheeran, 2006; James, 1890; Jeannerod, 1988; Lewin, 1926; Meyer & Kieras 1997; Prinz, 1987). Goals therefore may correspond to some cognitive entity that is active during the task episode (James, 1890; Kruglanski & Kopetz, 2009). The episode related entity may be the goal related program that controls and organizes cognition across time enabling relevant processing.

Possibility 3: What is hierarchically stored in memory, and captured through constructs like tasks and goals, may not be just propositional representations, but includes the episodic memories of task and goal execution (e.g. Jacoby & Brooks, 1984; Logan, 1988; Medin & Shaffer, 1978; Neill, 1997). Note that by episodic memory we mean memory of past behavioral and cognitive episodes; on the other hand, we call task episodes because of their temporally extended nature. It is possible that instances of task execution generate a whole host of episodic (explicit and implicit) memories (e.g. Jacoby & Brooks, 1984) organized around the goal (e.g. Logan, 1988), with re-execution strengthening the common elements across different iterations of the task episode and whittling away those that were specific to individual instances, and thus create episodic memories of task execution. Later execution of task episodes will occur through the recall and instantiation of the episodic memory of that task episode causing the signs of hierarchical cognition, whereby the episodic memory of the task episode may be assembled along with executive elements to create programs evidenced by the current study.

Possibility 4: It is well known that attention cannot be sustained indefinitely. Faced with an episodic task, cognition may instantiate attention in chunks that last for a time period and have to be re-instantiated. The instantiation of these chunks may be synced with conceived task boundaries. Hence, an attentional chunk may be instantiated at a beginning of a task-episode. The magnitude instantiated may be such that it lasts the estimated duration of the episode.

In other words, the subsuming program, instead of directly specifying the identity and sequence of component steps, instantiated goal related organizational changes, control or attention across time in order to ensure that correct component processes and actions are selected across time.

### Programs and Routines

Entities that are selected and instantiated as one unit but correspond to a sequence of processes and actions have been proposed and evidenced in other domains. Ullman (1984) suggested that the immediate and effortless perception of spatial properties and relations of objects may be achieved through automatic instantiation of a *visual routine* – a fixed set of elemental processes on early visual representations (for a related construct see *Sprites*, Cavanagh et al. 2001). Logan (1999) suggested that attention during perceptual tasks be considered as a routine (*attentional routine*) consisting of the set of processes that intervene between perception and reaching a goal relevant propositional conclusion, e.g. during identification of object X this routine would intervene between early perceptual processes and the reaching of conscious conclusion that the ‘visible object is X’. Component motor acts of a larger motor action come about through a common program assembled at the beginning of execution (Henry & Rogers, 1960; Keele, 1968; Schmidt, 1975). More recently, Botvinick, Niv and Barto (2009) proposed that extended sequence of behavior (e.g. ‘open laptop’, ‘mouse to browser icon’, ‘double-click’, ‘enter URL’, among others) may be chunked into larger routines (‘check email’) through hierarchical reinforcement learning wherein the reward signals strengthen the entire sequence of behavior, and not just one act, allowing instantiation of the entire sequence as a routine. The cognitive entity evidenced by task sequence studies (e.g. Schneider & Logan, 2006) may also be seen as programs that across time instantiate the task item related states (e.g. task rule related sets) that allow correct motor acts to be selected (see Discussion, Schneider & Logan, 2006).

When we execute our behavior we are aware of the task and the goal identities being executed (Vallacher & Wegner, 1989). These typically correspond to an episode and consist of numerous sequential acts (‘prepare breakfast’, ‘boil water’). Our phenomenal experience that we execute such task identities, and not individually execute its very many component acts, may have be real. To execute a task identity may be to execute the corresponding episode as one unit. Hence, cognition may prepare for the entire episode as one unit at the beginning. This may manifest as assembling a program – entities embodying this preparation. The nature of this program, and the prospective preparation it embodies, may depend on the familiarity and predictability of the task episode related to that goal. Completely predictable tasks may be prepared down to the level of individual motor acts to be made across time (e.g. Henry & Rogers, 1960; Rosenbaum et al. 1983). During less predictable task episodes like memorized task sequence execution component motor acts cannot be prepared for in advance. Here the program may involve preparing for the sequence of stimulus – response rules to be applied across time (e.g. Schneider & Logan, 2006). During unpredictable task episodes, the program may instantiate control and related goal-directed changes in cognition. These may set the top-down template for the search for the correct next step. Programs, therefore, may not just be limited to predictable tasks but may be found whenever an extended period of purposive cognition is given a distinct task and goal identity.

### Conclusions

We showed that signs of hierarchical cognition, identical to that seen previously during the execution of memorized and predictable behavioral sequence, can be seen whenever any extended arbitrary segment of behavior is construed as one task, even when the components of that extended behavior are unknown and unpredictable. These suggest that even unpredictable task episodes may be executed through subsuming higher-level episode related cognitive entities assembled at the beginning of execution. We argue that these entities may be seen as programs that are the means of organizing cognition across time into goal directed episodes.

